# Tipifarnib potentiates the antitumor effects of PI3Kα inhibition in *PIK3CA*- and *HRAS*-dysregulated HNSCC via convergent inhibition of mTOR activity

**DOI:** 10.1101/2023.01.17.523964

**Authors:** Alison E. Smith, Stacia Chan, Zhiyong Wang, Asako McCloskey, Quinn Reilly, Jayden Z. Wang, Hetika Vora Patel, Keiichi Koshizuka, Harris S. Soifer, Linda Kessler, Ashley Dayoub, Victoria Villaflor, Douglas Adkins, Justine Bruce, Alan Ho, Cesar Perez Batista, Glenn Hanna, Amaya Gascó Hernández, Andrew Saunders, Stephen Dale, J. Silvio Gutkind, Francis Burrows, Shivani Malik

## Abstract

Outcomes for patients with recurrent/metastatic (R/M) head and neck squamous cell carcinoma (HNSCC) are poor, with median overall survival ranging from 6 to 18 months. For those who progress on standard of care (chemo)immunotherapy, treatment options are limited, necessitating the development of rational therapeutic strategies. Toward this end, we targeted the key HNSCC drivers PI3K-mTOR and HRAS via the combination of tipifarnib, a farnesyltransferase inhibitor, and alpelisib, a PI3Kα inhibitor, in multiple molecularly defined subsets of HNSCC. We find that tipifarnib synergizes with alpelisib at the level of mTOR in PI3Kα-or HRAS-dependent HNSCCs, leading to marked cytotoxicity *in vitro* and tumor regression *in vivo*. Based on these findings, we have launched the KURRENT-HN trial to evaluate the effectiveness of this combination in PIK3CA-mutant/amplified and/or HRAS-overexpressing R/M HNSCC. Preliminary evidence supports the clinical activity of this molecular biomarker-driven combination therapy.

**Significance:** Backed by strong mechanistic rationale, the combination of alpelisib and tipifarnib has the potential to benefit >45% of R/M HNSCC patients. By blocking feedback reactivation of mTORC1, tipifarnib may prevent adaptive resistance to additional targeted therapies, thereby enhancing their clinical utility.

## Introduction

With nearly 900,000 new cases diagnosed each year, head and neck squamous cell carcinoma (HNSCC) is the sixth most prevalent cancer worldwide, and its incidence is increasing (*1*). HNSCC is a heterogeneous disease with tumors arising from epithelial linings of the oral cavity, pharynx, and larynx. Regardless of the subsite, locoregional recurrences and distant metastases are not uncommon despite curative-intent treatment with surgery, radiation, and/or chemotherapy. Recurrent/metastatic (R/M) HNSCC is a challenging disease to treat, and patient prognosis is poor, with a dismal median overall survival rate of about 12 months (*2*). In 2019, the immune checkpoint inhibitor pembrolizumab targeting PD-1 was approved as first line therapy for R/M HNSCC patients based on the findings of the KEYNOTE-048 trial (*3*). Patients treated with pembrolizumab plus chemotherapy (platinum + 5-fluorouracil, 5-FU) had an improved median overall survival of 13.0 months compared to 10.7 months for patients receiving cetuximab plus chemotherapy (HR 0.77; CI: 0.63-0.93; p = 0.0034). While pembrolizumab improved the duration of response in the monotherapy arm, the overall response rates in this study were the same as achieved by historical standard-of-care chemotherapy (platinum +5-FU +cetuximab) since 2008 (36%). Few options currently exist for patients who progress on (chemo)immunotherapy and to date, no biomarker-selected therapies targeted to molecular subtypes of head and neck cancers have been approved. Therefore, it is of critical importance to identify new therapies tailored to the specific molecular characteristics of HNSCCs.

In head and neck cancers, the PI3K-AKT-mTOR pathway is the most frequently dysregulated signaling cascade. Activation of this oncogenic signaling pathway is commonly achieved via amplification or gain-of-function mutation of *PIK3CA*, the gene encoding the α isoform of the PI3K p110 catalytic subunit, making PI3Kα an attractive therapeutic target in this setting (*4, 5*). Although the PI3Kα inhibitor alpelisib (Novartis) has shown some activity in HNSCC in early clinical trials (*6*), its efficacy as a single agent is limited by feedback reactivation of PI3K (*7-10*) or compensatory parallel pathways such as RAS-MAPK (*11-13*), necessitating the development of rational combination strategies. While combined EGFR and PI3K inhibition targets the signaling pathways of interest in HNSCC, significant toxicity was observed when alpelisib was combined with cetuximab (*14*). Thus, the mechanistic strategy must be weighed carefully against the toxicity profile of potential combination drugs. For example, though the combination of alpelisib with a rapalog or TOR kinase inhibitor would sustainably block pathway signaling, overlapping toxicities primarily attributed to mTORC2 inhibition make the combination hard to justify from a tolerability standpoint. An ideal combination partner would target the feedback-reactivated pathway(s) to synergize mechanistically with alpelisib, while maintaining acceptable tolerability.

Tipifarnib, a potent and selective farnesyltransferase inhibitor (FTI), can inhibit several proteins involved in oncogenic growth factor signaling, including HRAS and RHEB, which depend on farnesylation for their activation and/or cellular localization (*15, 16*). Tipifarnib’s ability to inhibit both HRAS and RHEB makes it a promising candidate partner drug for alpelisib in HNSCC. HRAS is the dominant RAS isoform in squamous cell carcinomas (*17*) and although all RAS isoforms regulate PI3K (*18, 19*), HRAS preferentially signals through this route (*20*). As RHEB is a non-redundant TORC1 activator (*21*), tipifarnib simultaneously acts as an inhibitor of mTOR, which has been shown to be an important driver of both tumor growth and PI3K inhibitor (PI3Ki) resistance in HNSCC (*22-24*). Indeed, sustained inhibition of mTORC1 activity is required for PI3Ki efficacy (*25, 26*). By blocking both HRAS-MAPK and PI3K-AKT-mTOR signaling, tipifarnib can potentially blunt the feedback (re)activation of the two major pathways driving PI3Ki resistance and greatly enhance the efficacy of alpelisib in head and neck cancers. Importantly, tipifarnib is well tolerated in humans as evidenced by over two decades of safety and tolerability data and has shown encouraging clinical activity in *HRAS*-mutant squamous cell carcinomas (*27, 28*).

Here, we demonstrate that combined alpelisib and tipifarnib is efficacious in preclinical models of HNSCC and provide mechanistic rationale for advancing this combination into the clinic. Via concurrent inhibition of HRAS and RHEB, tipifarnib blocks the feedback reactivation of MAPK and mTOR following alpelisib exposure, leading to durable pathway inhibition and tumor cell death in both PI3Kα and HRAS-dysregulated models. The KURRENT-HN trial (NCT04997902) is underway to evaluate this combination in recurrent/metastatic HNSCCs and encouraging preliminary results have been observed.

## Results

### The combination of alpelisib and tipifarnib inhibits spheroid growth in *PIK3CA-* and *HRAS*-dysregulated HNSCC cell line models

HRAS-MAPK and PI3K-AKT-mTOR are highly interdependent pathways in squamous cell carcinomas. HRAS requires PIK3α to transform squamous epithelial cells (*29*), helical domain *PIK3CA* mutants must bind RAS for transformation (*30*), and overexpression of mutant or WT *HRAS* in a *PIK3CA*-mutant HNSCC cell line drives resistance to alpelisib (*23*). Based on such close crosstalk between HRAS and PI3Kα, we hypothesized that the combination of tipifarnib and alpelisib would be effective in head and neck tumors with high *HRAS* expression, *PIK3CA* amplification, or *PIK3CA* mutations. To ascertain what proportion of head and neck cancers might benefit from this combination, we evaluated rates of mutation and amplification of *PIK3CA* and *HRAS* in the Cancer Genome Atlas (TCGA) dataset. Approximately 30% of head and neck tumors harbored mutations or amplification of *PIK3CA* (Fig. 1A). As has been previously described (*27*), *HRAS* mutations were relatively rare, occurring in 4-8% of HNSCCs. However, the fact that *HRAS* mutations are restricted to only a handful of cancer subtypes, including HNSCC, implies that HRAS may play an important role in these tumors (*31*). Consistent with this notion, squamous cell carcinomas expressed elevated levels of *HRAS* mRNA compared to other tumor types in the TCGA PanCancer Atlas dataset, with up to 15% of head and neck tumors exhibiting high expression (defined as *HRAS* expression of 1 standard deviation above the mean of HNSCC tumors) (Fig. 1B). Taken together, we hypothesize that approximately 45% of all HNSCC patients could potentially benefit from the simultaneous inhibition of HRAS and PI3K-mTOR signaling by tipifarnib and alpelisib.

**Figure 1.**
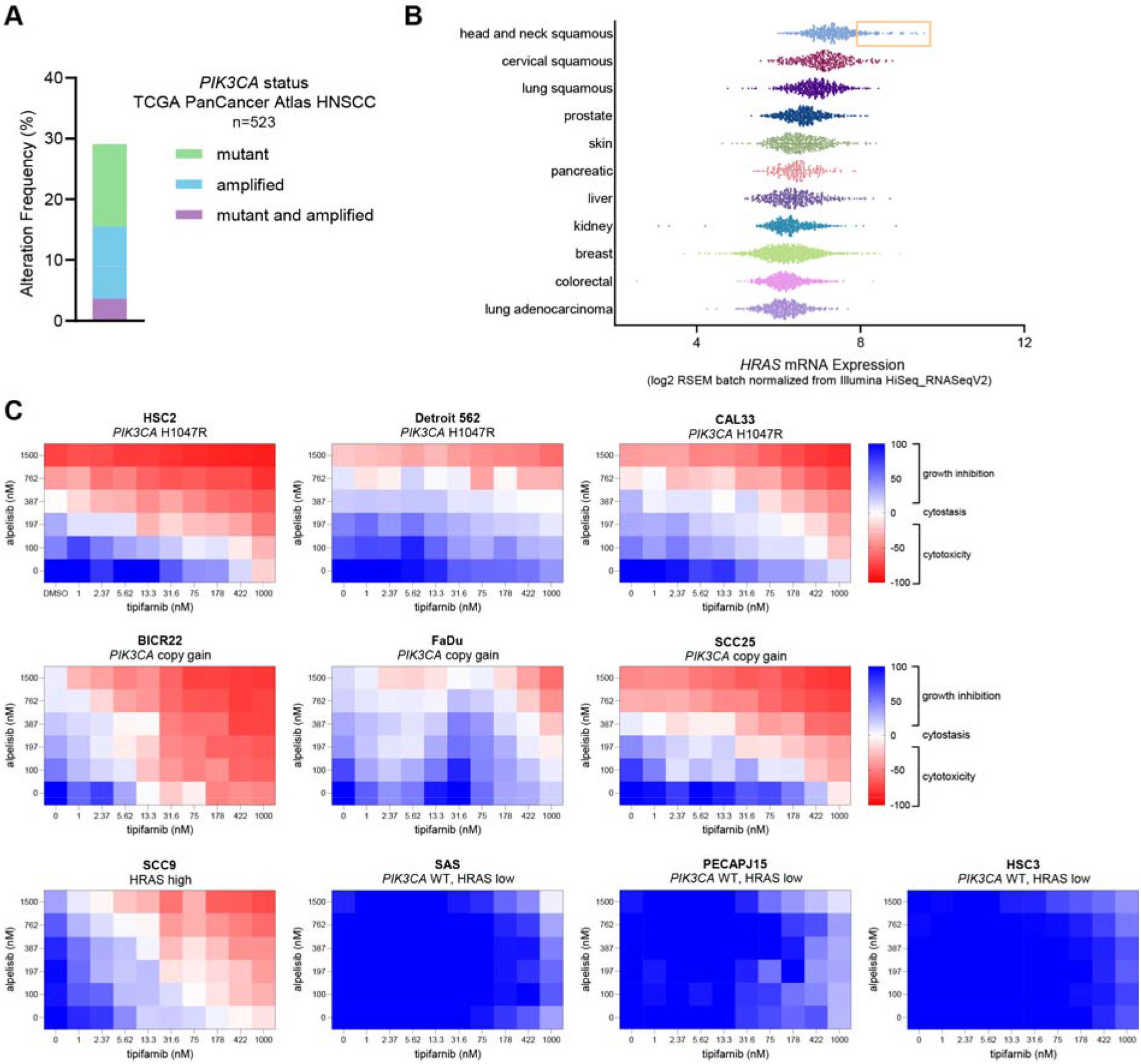
*PIK3CA* mutation/amplification and *HRAS* overexpression are common in HNSCC and cell lines harboring these alterations are sensitive to combined tipifarnib and alpelisib. **A**, Frequency of *PIK3CA* gain-of-function mutations and amplifications in the TCGA HNSCC dataset (n=523). **B**, *HRAS* mRNA expression in various tumor types in the TCGA PanCancer Atlas. Subset of HNSCCs with *HRAS* expression exceeding 1 standard deviation from the mean is highlighted by orange box. **C**, HNSCC cell lines of the indicated *PIK3CA/*HRAS statuses were cultured as 3D tumor spheroids and treated with increasing concentrations of tipifarnib and/or alpelisib for 7 days. Heat maps indicate the degree of growth inhibition or cytotoxicity induced by the compounds. Data are representative of 3 biological replicates.

To explore the therapeutic potential of this combination, we first assessed the sensitivity of a panel of *PIK3CA-* or *HRAS*-dysregulated HNSCC cell lines to tipifarnib and alpelisib. Cell lines with greater than 2.5 copies of *PIK3CA* were defined as copy gain and a cell line with the highest ratio of active (GTP-bound) to total HRAS was selected as the HRAS-high model (S1A, B). When cultured as 3D tumor spheroids and treated in a checkerboard dose-response fashion, cell lines with gain-of-function *PIK3CA* mutations, *PIK3CA* copy gain, or high HRAS activity were sensitive to combined tipifarnib and alpelisib (Fig. 1C). Additive or synergistic inhibitory effects on cellular viability were observed in these lines with induction or enhancement of cytotoxicity by the combination (Fig. S1C). In contrast, cell lines lacking both *PIK3CA* mutation/amplification and HRAS overexpression did not respond to either single agent or the combination (Fig. 1C). These results demonstrate that tipifarnib potentiates the antiproliferative effects of alpelisib in *in vitro* models of PI3Kα/HRAS-dysregulated HNSCC.

### Combined tipifarnib and alpelisib treatment inhibits mTOR and RSK more potently and durably than either agent alone

We next interrogated the molecular mechanism underlying the enhanced antiproliferative and cytotoxic activity of the combination compared to the single agents in our *in vitro* HNSCC models. As the single-agent efficacy of PI3K pathway inhibitors has historically been limited by feedback reactivation of PI3K-AKT-mTOR and RAS-MAPK signaling (*7-13*), we examined the effect of addition of tipifarnib on the kinetics of alpelisib’s inhibition of these pathways. In *PIK3CA* H1047R mutant CAL33 cells, alpelisib rapidly inhibited the activity of AKT, mTOR, and ERK, as indicated by dephosphorylation of their substrates PRAS40, S6K, and RSK, respectively, after 1 hour of treatment (Fig. 2A). However, by 24 hours, phosphorylation of these proteins partially or completely rebounded. In contrast, when cells were treated with tipifarnib (causing HRAS and RHEB to become defarnesylated, as indicated by mobility shift) 24 hours prior to addition of alpelisib, basal phosphorylation of S6K, S6, and RSK was reduced, the initial inhibition by alpelisib was deeper, and the subsequent rebound was blocked. This potent and durable inhibition of mTOR and RSK by the tipifarnib-alpelisib combination corresponded with cell cycle arrest (dephosphorylation of Rb) and induction of apoptosis (PARP, caspase 3, and caspase 7 cleavage) (Fig. 2A). Live cell imaging experiments corroborated these findings; combination treatment more potently induced cytotoxicity in CAL33 cells than tipifarnib or alpelisib alone (Fig. 2B). In *PIK3CA* WT, HRAS low HSC3 cells, neither PARP cleavage (Fig. S2A) nor Annexin V/cytotoxicity (Fig. S2B) were significantly induced by the combination.

**Figure 2.**
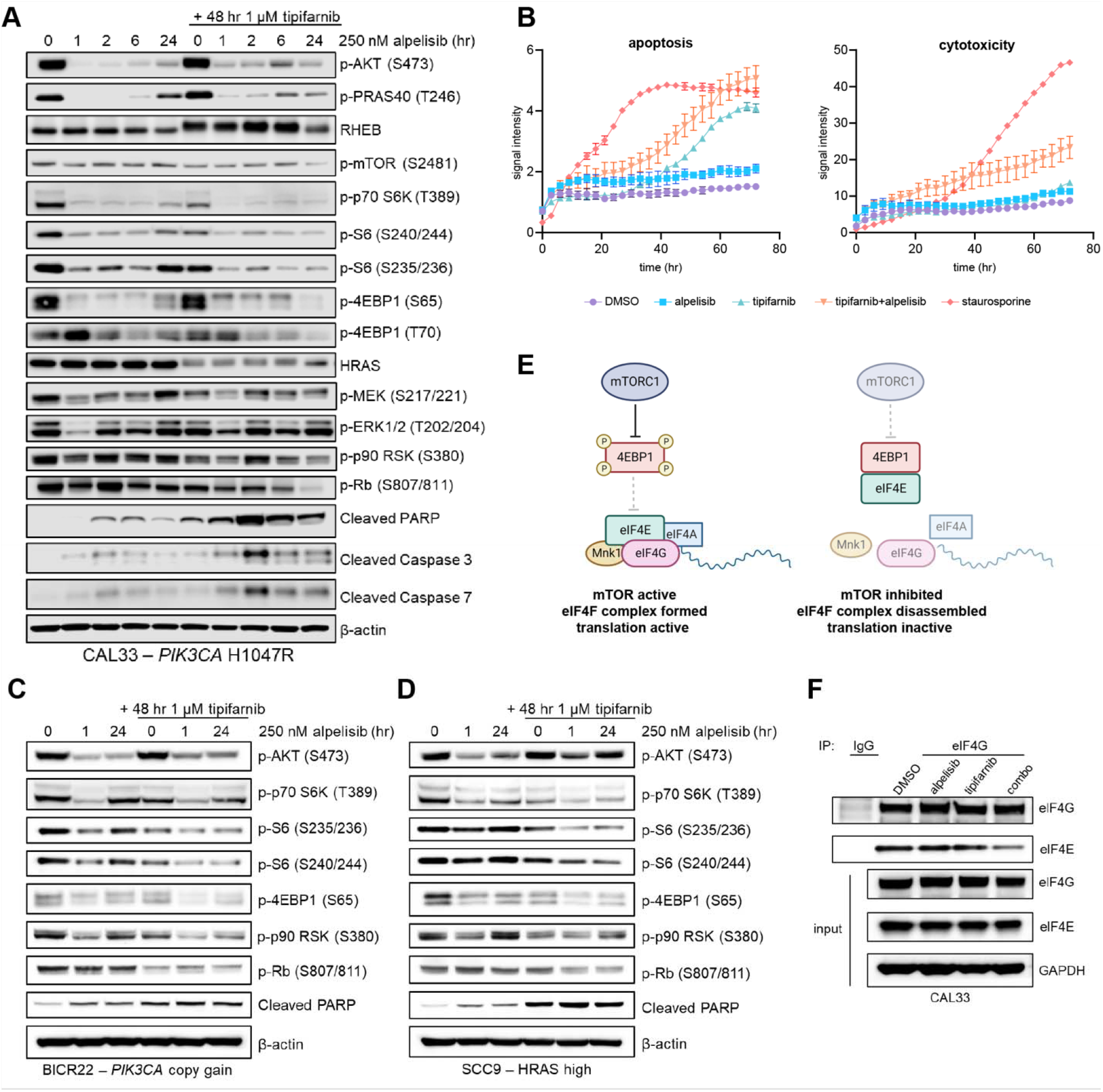
Tipifarnib blunts mTOR and RSK reactivation following alpelisib treatment and induces cell cycle arrest and apoptosis in *PIK3CA*-mutant, *PIK3CA*-amplified, and HRAS-high cell lines. **A**, Immunoblot of the indicated signaling proteins in *PIK3CA* mutant CAL33 cells treated with 250 nM alpelisib for 0, 1, 2, 6, or 24 hours in the presence or absence of 1 μM tipifarnib. Cells were treated with tipifarnib for 48 hours. Images are representative of 3 biological replicates. **B**, Apoptosis (Annexin V) and cytotoxicity (loss of membrane integrity, nuclei exposure) over time in CAL33 cells treated with DMSO, 250 nM alpelisib, 1 μM tipifarnib, or the combination for 72 hours measured via Incucyte live cell imaging. 100 nM staurosporine was used as a positive control. Data are means +/- SD of three biological replicates. **C and D**, Immunoblots of indicated proteins in **C**, *PIK3CA* copy gain BICR22 cells or **D**, HRAS-high SCC9 cells treated with 250 nM alpelisib for 0, 1, or 24 hours in the presence of absence of 1 μM tipifarnib (48-hour treatment). **E**, Graphical overview of role of mTORC1 in regulating assembly of the eIF4F translation initiation complex. mTORC1 phosphorylates 4EBP1, impeding its binding to eIF4E, allowing for complex assembly. **F**, eIF4G (or IgG control) was immunoprecipitated from CAL33 cells treated with DMSO, 250 nM alpelisib (24 hours), 1 μM tipifarnib (48 hours), or the combination. Immunoblots indicate the levels of eIF4G and eIF4E co-immunoprecipitated and in the input cell lysate. Images are representative of two biological replicates.

In *PIK3CA* copy gain BICR22 cells and HRAS-high SCC9 cells, combined tipifarnib-alpelisib treatment had a similar impact on signaling. Phosphorylation of S6K, S6, 4EBP1, and RSK was transiently reduced by alpelisib, but rebounded by 24 hours (Fig. 2C, 2D). Addition of tipifarnib reduced baseline phosphorylation of these proteins, deepened their inhibition by alpelisib, and blunted their rebound after 24 hours of alpelisib exposure. More potent TOR and RSK blockade correlated with diminished Rb phosphorylation and induction of PARP cleavage, indicating that the combination inhibited cell cycling and induced cell death, as in *PIK3CA*-mutant cells.

As we observed a synergistic inhibition of mTOR substrate phosphorylation by combined treatment with alpelisib and tipifarnib in our cell line models, we examined the effect these agents had on eIF4F complex formation. mTORC1 phosphorylates 4EBP1, preventing its binding to eIF4E, allowing eIF4E:eIF4G interaction and initiation of translation (Fig. 2E). In CAL33 cells, the interaction of eIF4E with eIF4G was markedly reduced by combination treatment, as measured by eIF4G immunoprecipitation (Fig. 2F). Thus, tipifarnib and alpelisib inhibit protein translation by targeting mTOR activity. Taken together, these findings demonstrate that addition of tipifarnib increases the depth and duration of mTOR inhibition compared to alpelisib alone, leading to cell death in *PIK3CA*/*HRAS*-dysregulated HNSCC.

### The combination of tipifarnib and alpelisib inhibits mTORC1 activity and induces tumor regression in a *PIK3CA*-mutant cell line-derived xenograft model

To determine whether enhanced sensitivity to the alpelisib-tipifarnib doublet also occurred *in vivo*, we implanted *PIK3CA-*mutant (CAL33) and *PIK3CA* WT/HRAS-low (HSC3) xenografts and treated the tumors with alpelisib, tipifarnib, or the combination. The combination blocked the growth of the *PIK3CA-*mutant tumors and induced tumor regression better than either agent alone (Fig. 3A). No additive effect was observed in the control *PIK3CA* WT/HRAS-low tumors (Fig. 3B).

**Figure 3.**
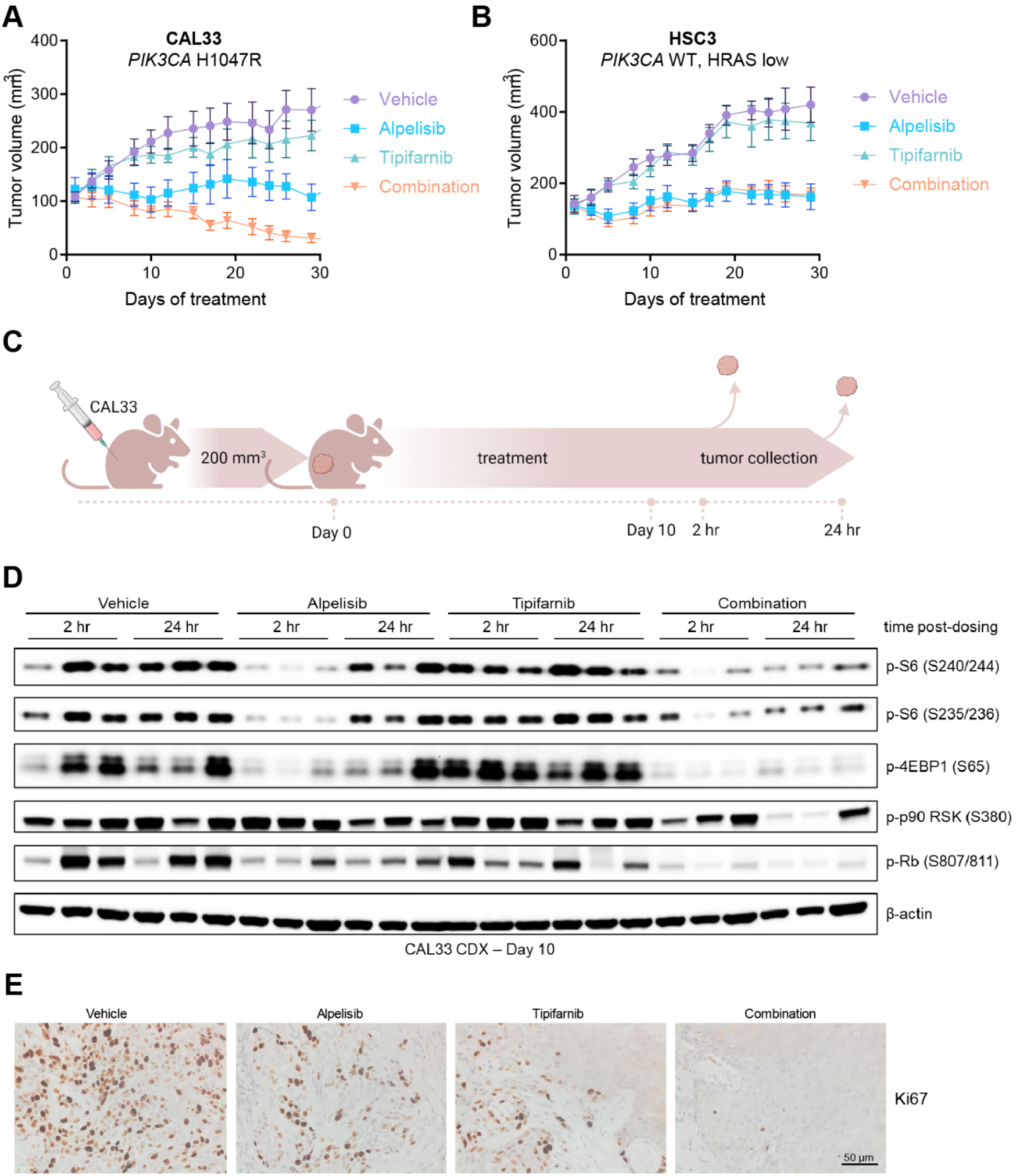
The tipifarnib and alpelisib doublet inhibits mTOR signaling and induces apoptosis and tumor regression in *PIK3CA* mutant cell line-derived xenograft tumors. **A and B**, Growth of **A**, CAL33 and **B**, HSC3 cell line-derived xenograft tumors treated with vehicle, tipifarnib (60 mg/kg BID), alpelisib (40 mg/kg QD), or the combination. Data represent means +/- SEM, n=10 mice per group. **C**. Overview of *in vivo* pharmacodynamic study. CAL33 cells were implanted in the flanks of athymic mice and allowed to reach 200 mm^3^, at which time treatment with vehicle, tipifarnib (60 mg/kg BID), alpelisib (40 mg/kg QD), or the combination was initiated and continued for ten days. On day 10, tumors were collected 2 hours and 24 hours after alpelisib dosing for analysis by immunohistochemistry and immunoblot. **D**, Immunoblots of indicated signaling proteins in CAL33 tumors treated with tipifarnib, alpelisib, or the combination for 10 days and collected at 2 and 24 hours post-alpelisib dosing. Tumors from three animals per treatment group/collection timepoint are shown. **E**, Representative immunohistochemical analysis of Ki67 in CAL33 tumors from panel **C** collected after 10 days of treatment, 24 hours after final alpelisib dose.

We next asked whether the mechanism underlying this tumor regression was consistent with that observed *in vitro*. We treated *PIK3CA*-mutant CAL33 tumors with tipifarnib, alpelisib, or the combination for ten days and collected tumors 2 and 24 hours after the final dose of alpelisib for immunohistochemistry and immunoblot analysis (Fig. 3C). Tipifarnib treatment resulted in defarnesylation of HRAS and RHEB (Fig. S3A). Alpelisib alone transiently inhibited mTOR activity (4EBP1, S6 phosphorylation at 2 hours), but by 24 hours, a significant rebound was observed (Fig. 3D). Addition of tipifarnib blocked this rebound and promoted cell cycle arrest as indicated by decreased Rb phosphorylation. Immunohistochemical analyses revealed the tipifarnib-alpelisib doublet inhibited tumor proliferation (Ki67 staining, Fig. 3E) and enhanced induction of apoptosis (PARP/caspase 3 cleavage, Fig. S3B) compared to monotherapy. Thus, the combination of alpelisib and tipifarnib durably inhibits mTORC1 activity *in vivo*, leading to cell death and tumor regression.

### Genetic depletion of RHEB and HRAS phenocopies tipifarnib treatment, implicating them as key farnesyltransferase targets in HNSCC

Although our mechanistic data point to HRAS and RHEB being the most important farnesylated targets in PI3Kα/HRAS-dysregulated HNSCCs, a number of signaling proteins are obligately farnesylated (*16*) and could contribute to tipifarnib sensitivity. Upon farnesyltransferase inhibition, these proteins are defarnesylated and lose their membrane localization and activity (Fig. 4A). Using subcellular fractionation, we confirmed that tipifarnib treatment resulted in the membrane delocalization of the majority of both HRAS and RHEB in CAL33, BICR22, and SCC9 cells (Fig. 4B-D). This result suggests that these proteins were sufficiently defarnesylated to impair their activity. Tipifarnib also displaced RHEB from lysosomes, a major site for mTORC1 activation (Fig. S4).

**Figure 4.**
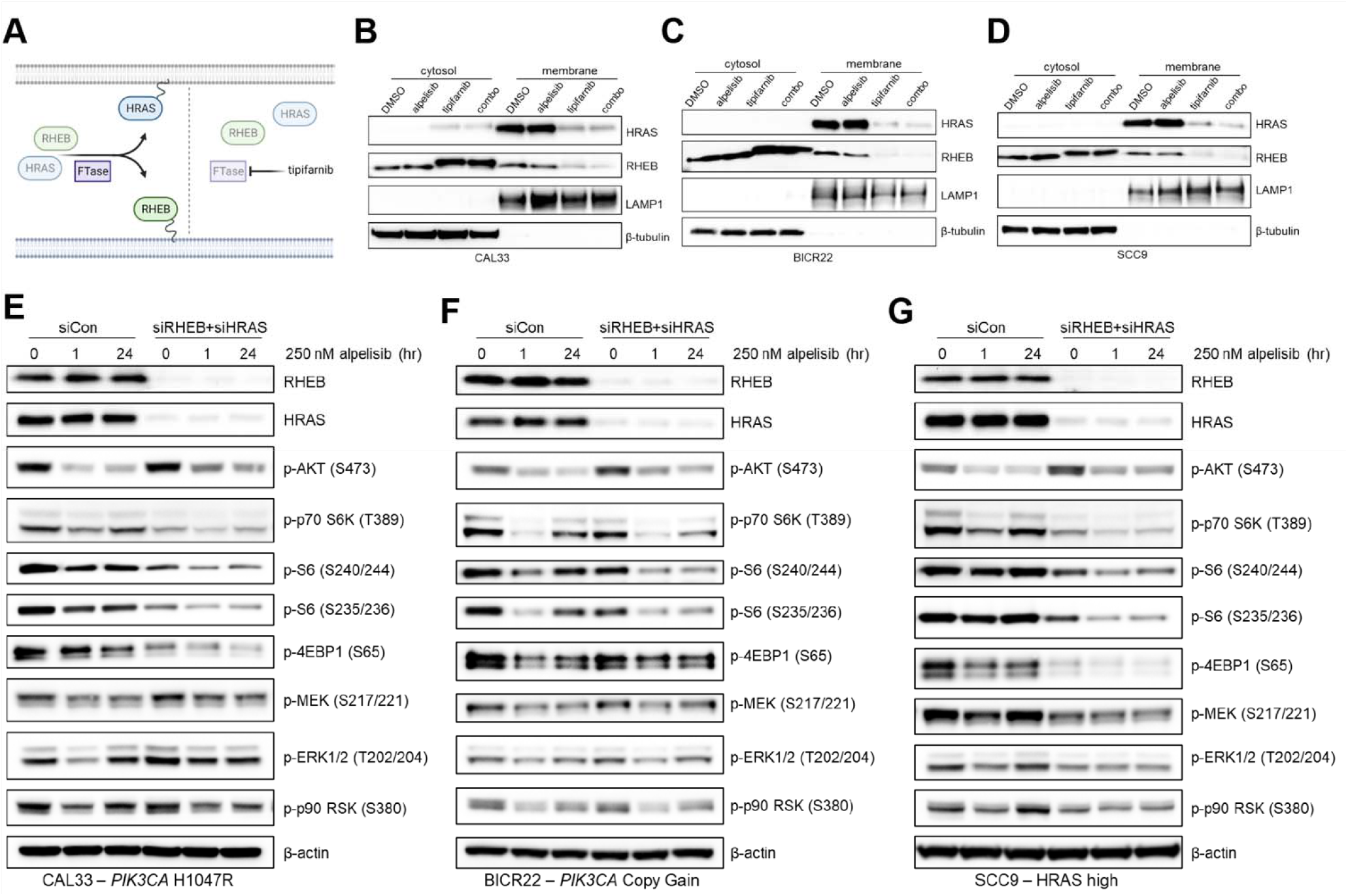
Depletion of the farnesyltransferase targets RHEB and HRAS phenocopies tipifarnib treatment in PIK3CA/HRAS dysregulated HNSCC cell lines. **A**. Graphical overview of the effect of tipifarnib on the membrane localization of HRAS and RHEB. Inhibition of farnesyltransferase (FTase) activity leads to defarnesylation of HRAS and RHEB and loss of membrane localization and activity. **B-D**. Immunoblots of HRAS, RHEB, LAMP1, and β-tubulin in the cytosol and membrane fractions of **B**, CAL33, **C**, BICR22, and **D**, SCC9 cells treated with DMSO, 250 nM alpelisib (24 hr), 1 μM tipifarnib (48 hr), or the combination. **E-G**. Immunoblots of indicated signaling proteins in **E**, CAL33, **F**, BICR22, and **G**, SCC9 cells treated with small interfering RNAs to knock down RHEB and HRAS expression (vs. control non-targeting pool) for 48 hours prior to addition of 250 nM alpelisib. Cells were collected and lysed after 0, 1, or 24 hours alpelisib treatment and immunoblot analysis performed.

To further clarify the role of RHEB and HRAS in mediating tipifarnib response, we asked whether genetic depletion of these two targets was sufficient to phenocopy the effect of tipifarnib on cell signaling. We transfected cells with small interfering RNAs to deplete *RHEB* and *HRAS* expression or a control non-targeting pool and, after allowing 48 hours for knock down, treated the cells with alpelisib. Like tipifarnib treatment, reduction of *RHEB* and *HRAS* expression decreased basal phosphorylation of S6K, S6, and 4EBP1 to varying degrees across the PIK3CA-mutant/amplified and high active cell lines (Fig. 4E-G). Inhibition of phospho-S6K, phospho-S6, and phospho-4EBP1 by alpelisib was greater in the double knock-down (siRHEB and siHRAS) cells compared to control-transfected cells, and this inhibition was more durable, exhibiting minimal rebound of phosphorylation after 24 hours. The effect of co-depletion of *RHEB* and *HRAS* expression on mTOR substrate phosphorylation was greatest in *PIK3CA*-mutant cells (Fig. 4E), whereas its effect on MAPK activity was largest in HRAS-high cells (Fig. 4G), pointing to a potential hierarchy of target dependence corresponding to genotype. As genetic targeting of HRAS and RHEB closely mimicked tipifarnib treatment in these models, we surmise these are the key farnesylation-dependent proteins in the context of PI3Kα inhibition in HNSCC.

### Combined, synchronous tipifarnib-alpelisib treatment robustly inhibits the growth of patient-derived xenograft models harboring *PIK3CA* alterations or HRAS overexpression

We next assessed the sensitivity of a panel of *PIK3CA-*mutant, *PIK3CA-*amplified, or HRAS-overexpressing (Fig. S5) patient-derived xenograft (PDX) models to single agent tipifarnib/alpelisib or the combination. Combination treatment consistently slowed or blocked tumor growth more effectively than alpelisib or tipifarnib alone, and induced tumor regressions in several models (Fig. 5A-C).

**Figure 5.**
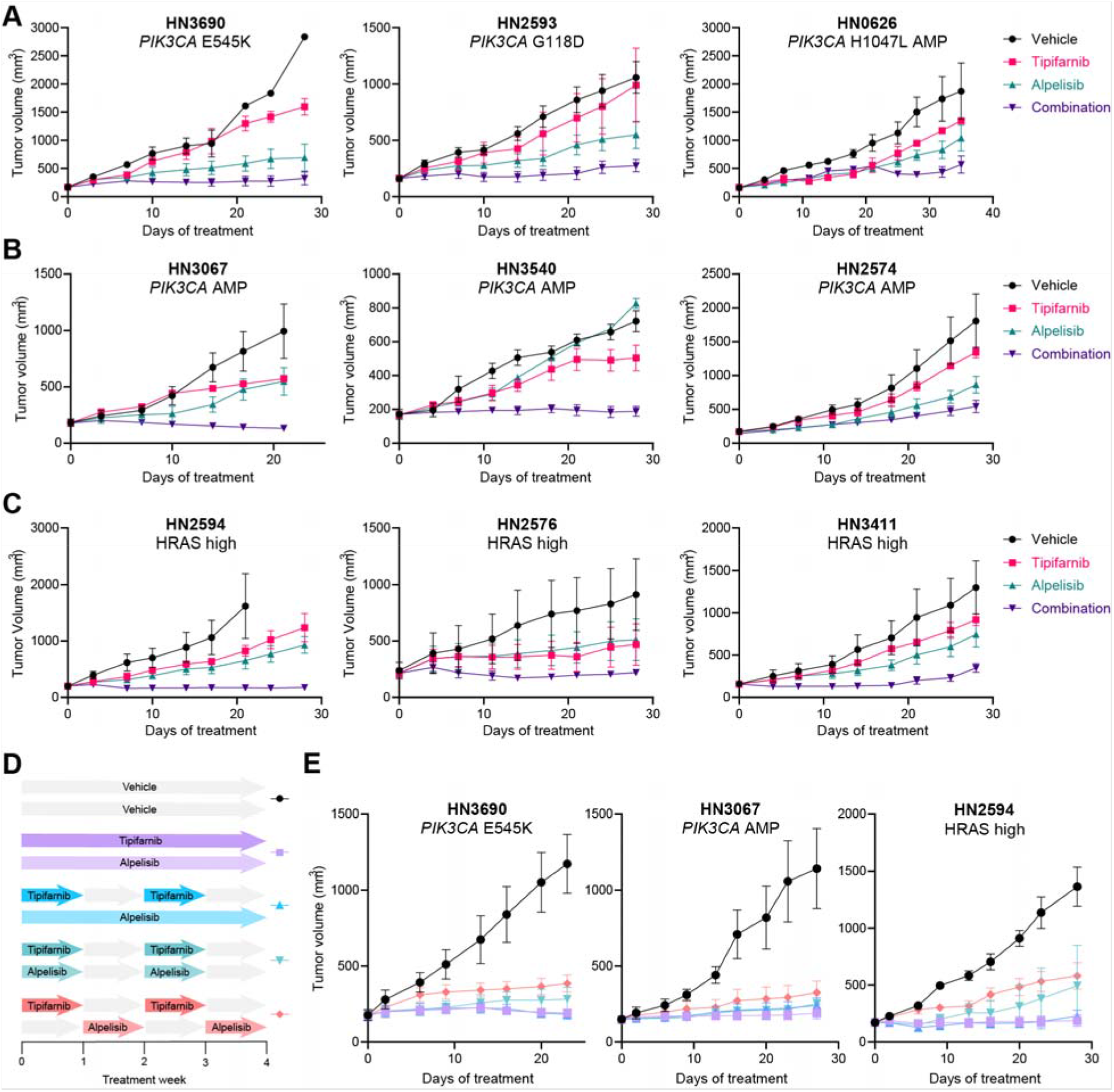
Combined, synchronous tipifarnib-alpelisib treatment robustly inhibits the growth of *PIK3CA*- and *HRAS*-dysregulated HNSCC patient-derived xenograft models. **A-C**, Growth of HNSCC PDX tumors harboring **A**, the indicated *PIK3CA* mutations, **B**, *PIK3CA* amplification (AMP), or **C**, *HRAS* overexpression (models in the top 25% of *HRAS* expression), treated with vehicle, tipifarnib (60mg/kg BID), alpelisib (40 mg/kg QD), or the combination on a continuous schedule. Data represent means +/- SEM, n=3 mice per group. **D**, Schematic overview of the tipifarnib (60 mg/kg BID) and alpelisib (40 mg/kg QD) dosing schedules evaluated in select HNSCC PDX models. **E**, Impact of the dosing schedules outlined in **D** on the growth of *PIK3CA-* or *HRAS*-dysregulated PDX tumors. Data represent means +/- SEM, n=5 mice per group.

The extensive crosstalk between the HRAS-MAPK and PI3K-AKT-mTOR pathways implies that synchronous administration of tipifarnib and alpelisib would have the greatest antitumor effects. However, concomitant blockade of multiple oncogenic signaling pathways can be challenging in the clinic. To explore scheduling options and the role of pathway interdependence in the observed combination activity, we tested various dosing schedules in *PIK3CA-* and HRAS-dysregulated PDX models (Fig. 5D). Continuous dosing of tipifarnib and alpelisib was most effective in blocking tumor growth but administering tipifarnib on alternate weeks (which is the established clinical regimen in HRAS-mutant HNSCC) while dosing alpelisib continuously was nearly as efficacious (Fig. 5E). Dosing alpelisib every other week slightly reduced the activity of the combination. Nevertheless, the observation that synchronous weekly on/off dosing retains robust activity may provide flexibility to optimize the dose scheduling of the combination in the clinic. It is important to note that non-synchronous intermittent dosing of alpelisib and tipifarnib was markedly less effective, highlighting the degree of cooperativity between HRAS/RHEB/PI3K and the synergistic potential of combination treatment. Although synchronous and non-synchronous discontinuous dosing appears similarly effective in the HRAS-high PDX model (Fig. 5E, right panel), this is due to a single outlier in the synchronous group. In this cohort, 4/5 tumors were maintained in full stasis with one rapidly growing breakout tumor, whereas 4/5 animals displayed tumor progression in the non-synchronous cohort.

### Preliminary evidence of clinical activity

Based on these findings, we launched the KURRENT-HN trial (NCT04997902) to assess the safety, tolerability, pharmacokinetics, pharmacodynamics, and preliminary antitumor activity of combined tipifarnib and alpelisib. This study is a phase 1/2 open label dose escalation study enrolling HNSCC patients with *PIK3CA* mutation/amplification and/or HRAS overexpression that have progressed on at least one prior line of therapy (Fig. 6A). While this clinical trial is ongoing, a 35-year-old male non-smoker with HPV16+ squamous cell carcinoma of the tonsil that had progressed following cisplatin/radiation and cemiplimab/ISA101b treatment was enrolled in the second dose cohort (Fig. 6A) based on a *PIK3CA* R88Q mutation in his tumor. The patient was administered tipifarnib and alpelisib and after one 28-day cycle, experienced an 81% reduction and a complete disappearance of the two target lesions in the lung. After three cycles, the complete response of the lung was maintained, and the patient had an 84% reduction in the target lesions. Since being on treatment, the patient has also experienced improvement in respiratory symptoms (Fig. 6B). As of the most recent data cut, the response was maintained, and the patient remained on study.

**Figure 6.**
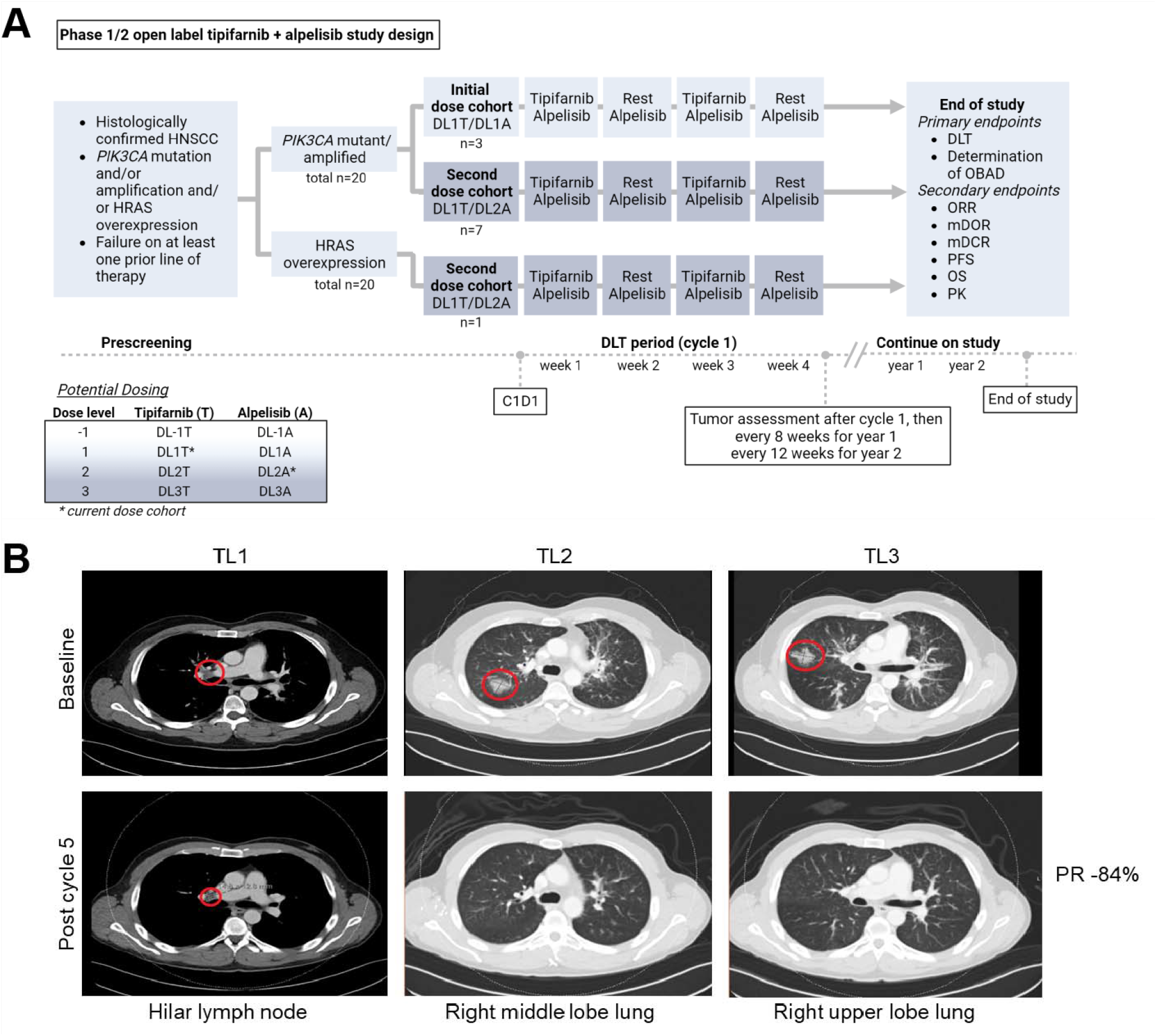
Clinical activity of combined tipifarnib and alpelisib in patient with advanced HNSCC. **A**, Overview of the KURRENT-HN trial (NCT04997902), enrolling patients with *PIK3CA-* mutant/amplified and/or HRAS-overexpressing HNSCCs. Dose escalation for tipifarnib and alpelisib determined by the Bayesian Logistic Regression Model in addition to clinical data reviewed by the safety monitoring committee. Abbreviations: DCR = disease control rate; DL = dose level; DLT = dose limiting toxicity; DOR = duration of response; OBAD = optimal biologically active dose, ORR = overall response rate; OS = overall survival; PFS = progression free survival; PK = pharmacokinetics **B**, Computed tomography scans of patient with *PIK3CA*-mutant metastatic HNSCC. Target lesions (red circles) are in the hilar lymph node and right upper lobe of the lungs. Patient experienced a partial response (PR) and remains on study.

## Discussion

Improvement in outcomes for patients with recurrent, metastatic head and neck squamous carcinoma has been hindered by a lack of targeted therapies. In this study, we provide *in vitro* and *in vivo* evidence that the mechanistically designed, biomarker-matched strategy of combining alpelisib and tipifarnib may be efficacious in *PIK3CA-* and *HRAS*-dysregulated HNSCCs. Showing encouraging early activity in the clinic and a manageable safety profile (NCT04997902), this combination has potential to benefit approximately half of HNSCC patients, for whom treatment options are currently limited.

As approximately 30% of head and neck cancers harbor alterations in *PIK3CA*, pharmacological targeting of p110α is an appealing therapeutic strategy. Unfortunately, feedback reactivation of PI3K and/or its parallel pathways greatly reduces the single-agent effectiveness of PI3K inhibitors. For example, in estrogen receptor-positive (ER+) breast cancer, where alpelisib is approved in combination with the ER antagonist fulvestrant (*32*), PI3Kα inhibition induces ER signaling while the latter activates the PI3K pathway as a mechanism of endocrine therapy escape (*33-35*). When combined, alpelisib and fulvestrant effectively block the bypass mechanisms tumor cells employ to evade the inhibitory effects of alpelisib monotherapy, leading to superior anti-tumor activity.

In HNSCC, mTOR and RAS-MAPK are the primary pathways driving adaptive resistance to PI3K inhibitors, but co-targeting PI3K and these escape pathways has proven challenging in the clinic. Direct inhibition of PI3K and mTOR via dual kinase inhibitors demonstrated unacceptable toxicity and/or lacked meaningful clinical activity (*36, 37*). Although the combination of alpelisib and the allosteric mTOR inhibitor everolimus was tolerable in solid tumors (like breast cancer and renal cell carcinoma), overlapping toxicities resulting from PI3K-mTOR pathway inhibition, such as hyperglycemia and stomatitis, make it difficult to assess the overall benefit-risk ratio (*38, 39*). Similarly, while backed by a strong mechanistic rationale, the clinical development of therapeutic strategies combining inhibitors of the PI3K and MAPK pathways has been hindered by safety concerns. The combination of PI3K and MEK inhibition showed favorable activity in solid cancers but at the expense of tolerability (*40*). Thus, further effort is needed to design translatable methods of co-inhibiting the PI3K-mTOR and RAS-MAPK pathways.

In this work, we propose a new therapeutic strategy—the combination of the farnesyltransferase inhibitor tipifarnib with alpelisib—to target the crosstalk of the PI3K and MAPK pathways in HNSCCs and limit their feedback reactivation upon PI3K inhibition. We demonstrate that tipifarnib achieves both vertical and lateral inhibition of PI3K signaling via depletion of RHEB and HRAS. Treatment of PI3Kα- and HRAS-dysregulated HNSCC cells with alpelisib exposes a dependency on mTOR that is effectively blocked by tipifarnib, thereby greatly enhancing sensitivity to PI3Kα inhibition.

Importantly, tipifarnib specifically inhibits mTORC1 (via inhibition of RHEB farnesylation), the driver of feedback-mediated resistance to alpelisib, while sparing mTORC2, the primary source of TOR inhibitor-associated toxicity (*41, 42*). Tipifarnib blocks localization of RHEB to all cellular membranes, including the lysosomes, a key site for RHEB’s regulation of mTORC1 activity. Recently, a small pool of RHEB was reported to be functional in the nucleus and found to be refractory to farnesyl transferase inhibition (*43*). While the exact role of this nuclear form of RHEB is unclear, exclusive inhibition of membrane-bound RHEB by tipifarnib may be a desirable property from a tolerability perspective when combining with a PI3K inhibitor.

Though RHEB has been nominated, albeit infrequently, as an oncogenic driver and therapeutic target in cancer (*44-47*), this work is the first to demonstrate that RHEB inhibition can prevent adaptive resistance to a targeted therapy. We describe how RHEB-mTORC1 blockade via tipifarnib potentiates the antitumor effects of PI3K inhibition in HNSCC, but sufficient TORC1 suppression is a common requirement for sensitivity to numerous other targeted agents, including inhibitors of RAF (*48*) and KRAS G12C (*49*). Positioned downstream of the majority of oncogenic drivers, mTORC1 is a critical mediator of tumor growth, survival, and therapy escape. We posit that by blocking feedback reactivation of mTORC1, tipifarnib may enhance the efficacy and expand the utility of other targeted therapies approved or currently in development, broadly benefiting patients with advanced malignancies.

## Methods

### Cell lines and reagents

Cell lines were obtained from ATCC (SCC25, SCC9, FaDu, and Detroit 562), JCRB (SAS, HSC2, HSC3), DSMZ (CAL33), or Sigma (PECAP15J) and maintained in a humidified atmosphere with 5% CO_2_ at 37°C. Cells were maintained in DMEM (Detroit 562, FaDu, HSC2, HSC3, CAL33, PECAP15J) or DMEM/F12 (SCC25, SCC9, SAS) supplemented with 10% FBS and penicillin/streptomycin. All lines tested negative for mycoplasma. Alpelisib was purchased from MedChemExpress and dissolved in DMSO. Dharmacon ON-TARGETplus siRNA SMARTPools against *RHEB* and *HRAS* were obtained from Horizon Discovery and transfected using Lipofectamine RNAiMAX (ThermoFisher).

### Tumor spheroid growth assays

Cells were resuspended in 4% Matrigel and seeded in 96-well ultralow attachment plates at a density of 1-1.5K cells/well. The following day, spheroids were treated with alpelisib and/or tipifarnib and baseline growth measured using 3D Cell Titer Glo reagent (Promega). Spheroids were incubated with drug for 7 days and a final CTG reading taken. Percentage growth was calculated by[(Ti-Tz)/(C-Tz)] x 100 for concentrations for which Ti>/=Tz and [(Ti-Tz)/Tz] x 100 for concentrations for which Ti<Tz, where Tz=time zero, C=control growth, and Ti=test growth at each drug concentration.

### Xenograft models and in vivo models

Cell line-derived xenograft experiments were performed at the University of California, San Diego (San Diego, CA), under protocol ASP # S15195, approved by the Institutional Animal Care and Use Committee (IACUC).

Patient-derived xenograft experiments were conducted at Crown Bioscience Beijing. The protocol and any amendment(s) or procedures involving the care and use of animals were approved by the IACUC of Crown Bioscience prior to initiation. During the study, the care and use of animals was conducted in accordance with the regulations of the Association for Assessment and Accreditation of Laboratory Animal Care.

### Immunoblotting

Cell lysates were prepared on ice by washing cells once with PBS, resuspending in 1X cell lysis buffer (Cell Signaling Technology #9803) or RIPA buffer supplemented with Halt protease inhibitor cocktail (Thermo Scientific #78430) and briefly sonicating or vortexing. Lysates were cleared by centrifugation (maximum speed, 10min) and protein concentration determined by BCA assay (Pierce). 20-60 ug of lysate was loaded on to 4-12% Bis-Tris gels (NuPAGE, Invitrogen) for electrophoresis and immunoblotting. Active RAS was detected using Thermo Scientific kit #16117. Membrane and cytosolic fractions were isolated using the Mem-PER Plus Membrane Protein Extraction Kit (Thermo Scientific kit #89842).

## Supplementary data

**Figure S1.**
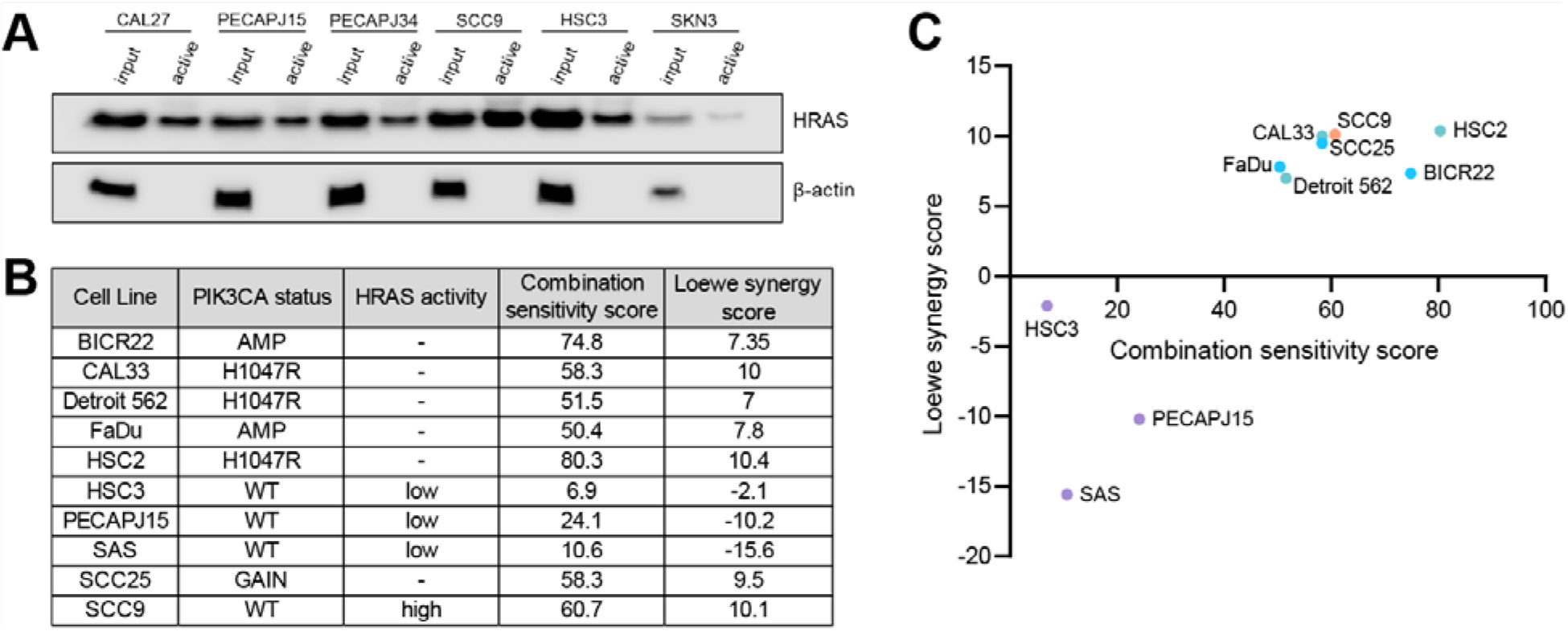
HNSCC cell line panel utilized to evaluate tipifarnib-alpelisib combination *in vitro*. **A**, Immunoblot of active and total HRAS in *PIK3CA* WT cell lines. Active/GTP-bound HRAS was assessed by pull-down with Raf1 RAS-binding domain. **B and C**, *PIK3CA* and *HRAS* status of cell lines evaluated for tipifarnib-alpelisib response in this study, and quantification of their combination sensitivity/synergy using the SynergyFinder R package (*50*). Loewe synergy score of less than -10 indicates antagonistic combination activity, -10 to 10 an additive effect, and greater than +10 a synergistic effect. SCC9 line harbors *HRAS* L133R mutation which does not lie within any functional domains of HRAS and therefore was determined not to be activating.

**Figure S2.**
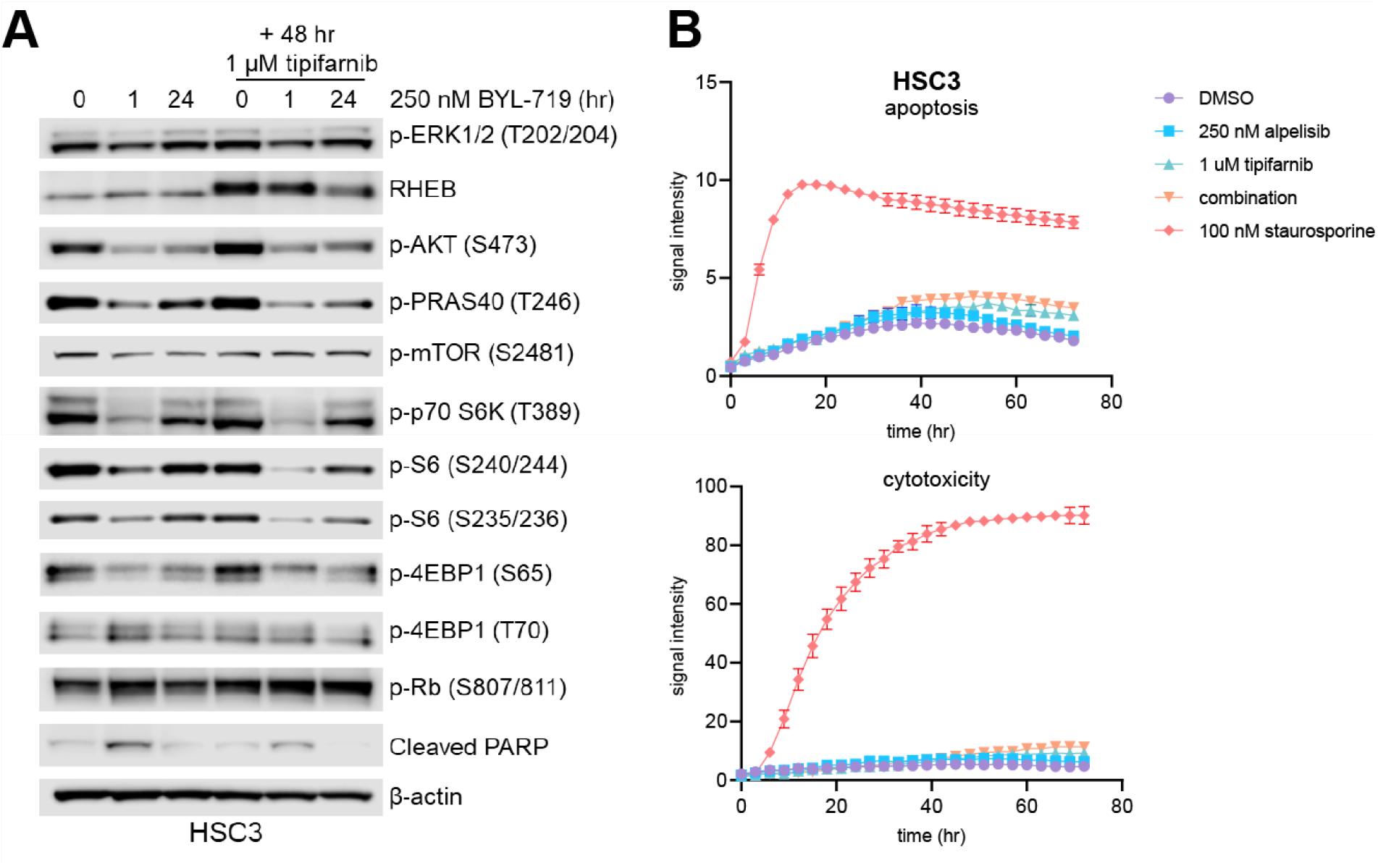
Cell line model without *PIK3CA* alteration or elevated HRAS activity is insensitive to the tipifarnib-alpelisib combination. **A**, Immunoblot of the indicated signaling proteins in control (WT *PIK3CA*/*HRAS*) HSC3 cells treated with 250 nM alpelisib for 0, 1, or 24 hours in the presence or absence of 1 μM tipifarnib. Cells were treated with tipifarnib for 48 hours. Images are representative of 3 biological replicates. **B**, Apoptosis (Annexin V) and cytotoxicity (DNA stain/loss of membrane integrity) over time in HSC3 cells treated with DMSO, 250 nM alpelisib, 1 μM tipifarnib, or the combination for 72 hours measured via Incucyte live cell imaging. 100 nM staurosporine was used as a positive control. Data represent means +/- SD of three biological replicates.

**Figure S3.**
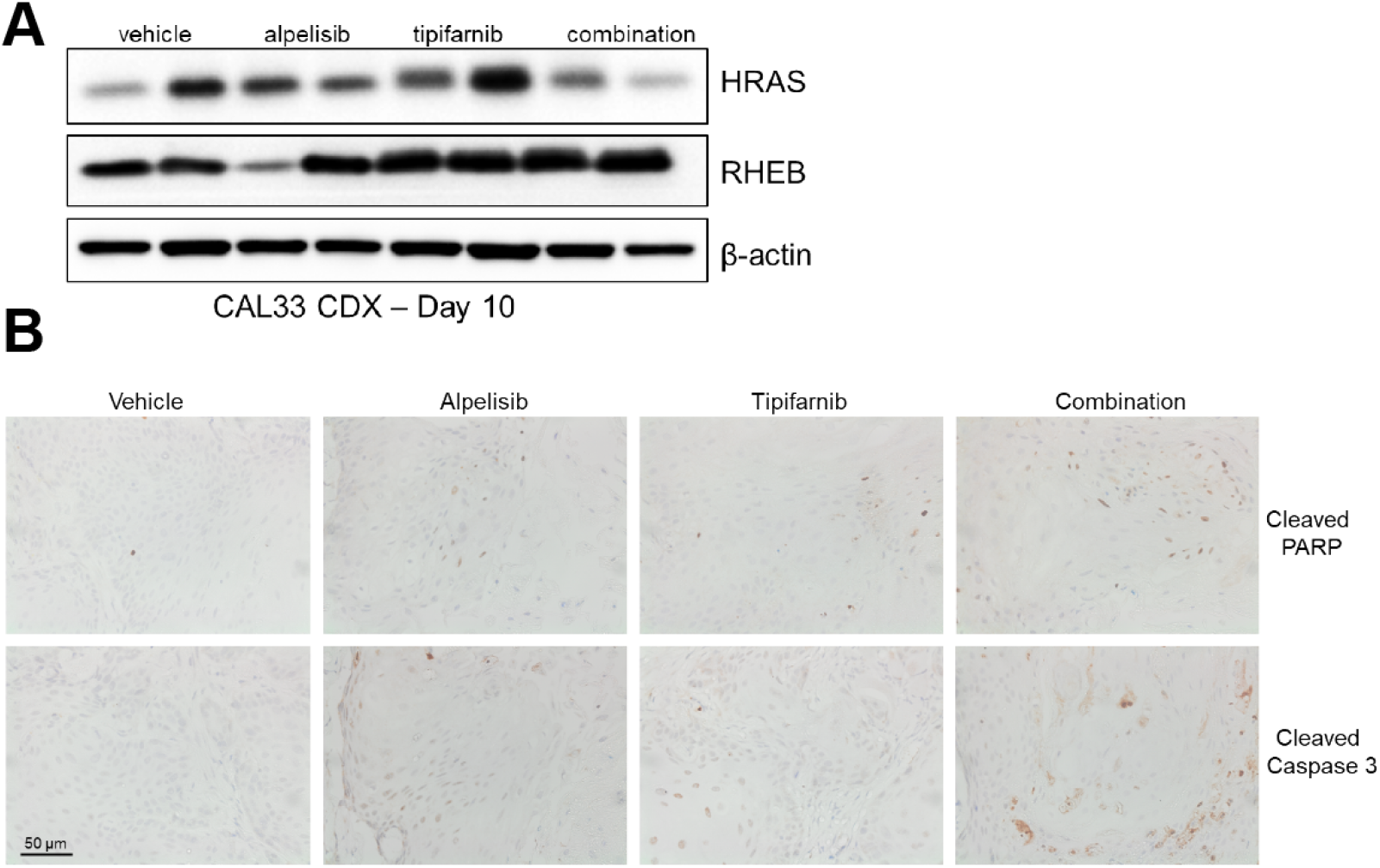
Activity of tipifarnib and alpelisib *in vivo*. **A**, Immunoblots of HRAS, RHEB, and beta-actin in CAL33 xenograft tumors treated with vehicle, alpelisib, tipifarnib, or the combination for 10 days. Two tumors per treatment group are shown. **B**, Representative immunohistochemical analysis of cleaved PARP and caspase 3 in CAL33 tumors exposed to indicated treatments for 10 days.

**Figure S4.**
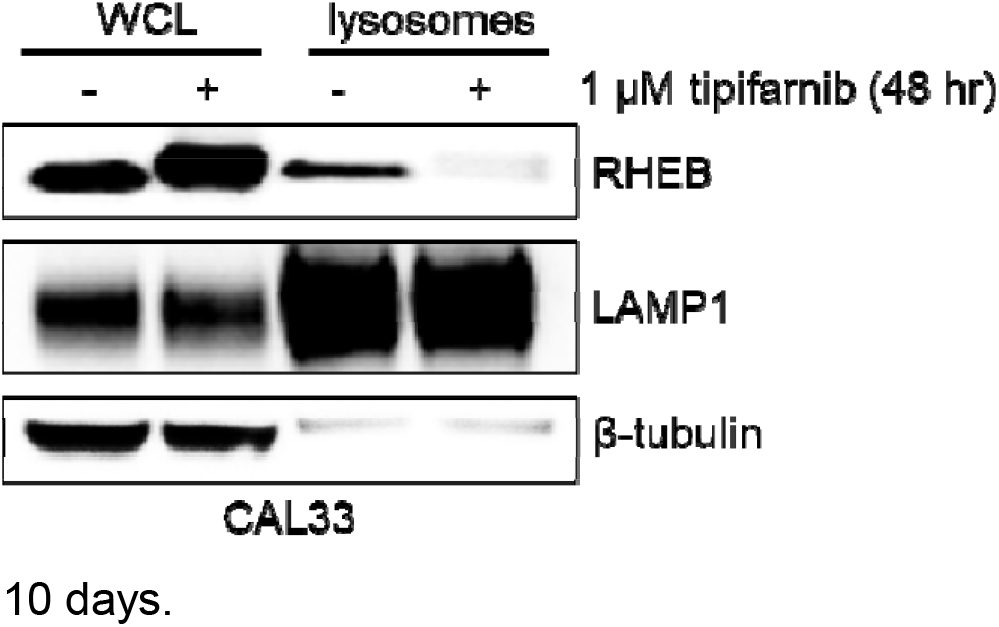
Tipifarnib blocks RHEB’s localization to lysosomes. Density gradient ultracentrifugation was utilized to extract lysosomes from CAL33 cells treated with DMSO or tipifarnib. Lysosomes were lysed and subjected to immunoblot analysis alongside whole-cell lysate (WCL). LAMP1 is a lysosome-specific marker.

**Figure S5.**
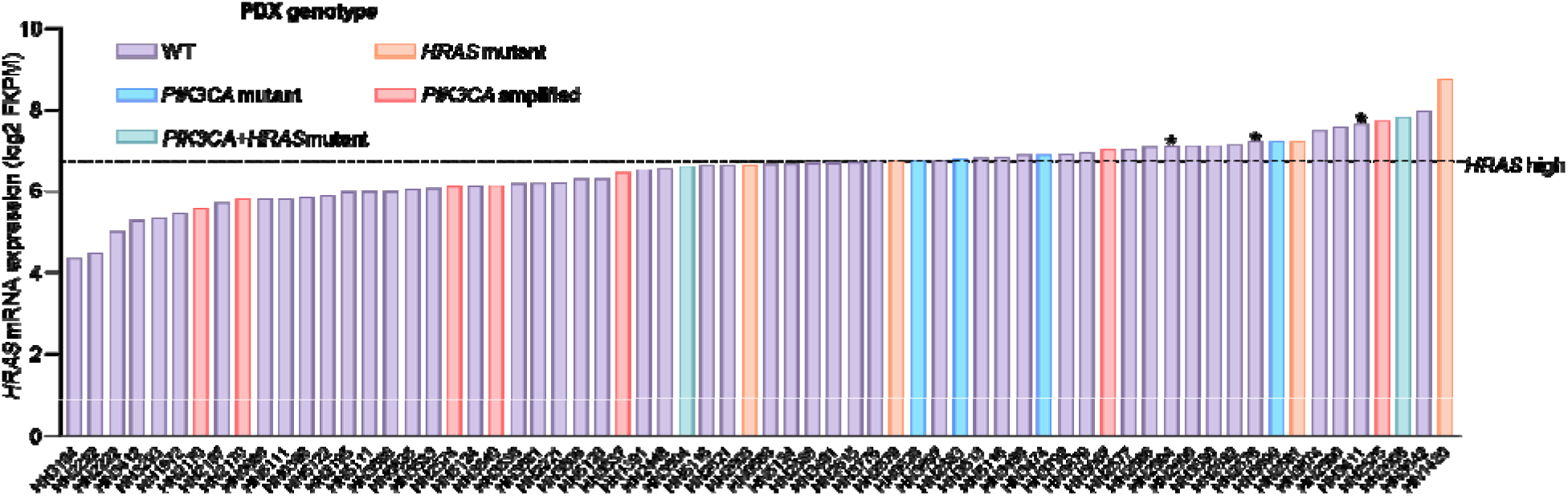
*HRAS* mRNA expression across Crown Bioscience’s HNSCC PDX models. Three models in the top 25% of expression, indicated by asterisks, were selected as HRAS-high models evaluated for sensitivity to tipifarnib and alpelisib in Fig. 5.

